# Canine tumor mutation rate is positively correlated with TP53 mutation across cancer types and breeds

**DOI:** 10.1101/2020.07.15.205286

**Authors:** Burair A. Alsaihati, Kun-Lin Ho, Joshua Watson, Yuan Feng, Tianfang Wang, Shaying Zhao

## Abstract

Spontaneous canine cancers are a valuable but relatively understudied and underutilized model in cancer research. To enhance their usage, we reanalyzed whole exome sequencing data published for 601 dogs with mammary cancer, osteosarcoma, oral melanoma, lymphoma, glioma or hemangiosarcoma from over 35 breeds, after rigorous quality control, including breed validation. Each cancer type harbors distinct molecular features, with major pathway alterations matching its human counterpart (e.g., PI3K for mammary cancer and p53 for osteosarcoma). On average, mammary cancer and glioma have lower mutation rates (median <0.5 mutation per Mb), whereas oral melanoma, osteosarcoma and hemangiosarcoma have higher mutation rates (median ≥1 mutation per Mb). Across cancer types and across breeds, the mutation rate is strongly associated with *TP53* mutation, but not with *PIK3CA* mutation. The mutation rate is also associated with a mutation signature enriched in osteosarcoma of Golden Retrievers, independent of *TP53* mutation. Finally, compared to other breeds examined, DNA repair genes appear to be less conserved in Golden Retriever which is predisposed to numerous cancers.

## Introduction

Cancers in pet dogs occur spontaneously in animals with an intact immune system, which gives them an advantage over traditional cancer models such as cell lines and rodents. These canine cancers more accurately emulate human cancers in etiology, complexity, heterogeneity, behavior, treatment and outcome. These similarities give them the potential to effectively bridge a gap between preclinical models and human clinical trials, accelerating bench-to-bedside translation, and as such, the National Cancer Institute (NCI) has recently issued programs targeting canine cancers. These include funding multi-institute immunotherapy trials with pet dogs and a 5-year project to build the NCI Integrated Canine Data Commons, a database to disseminate canine molecular and clinical data to the public.

However, current deficiencies create roadblocks to the effective use of canine cancers. This is clearly exemplified by sequence mutation, a hallmark of cancer^1^. Tumor mutation rate, mutation signature and mechanism are extensively investigated for nearly every type of human cancer via pan-cancer studies^2–5^. However, to our knowledge, no pan-cancer research has been published for the dog, and fundamental questions remain unanswered. For example, mutation rate varies significantly among human cancer types^6,7,8{Alexandrov, 2020 #1546^, and mutation rate-based subtypes have been identified in multiple cancers in human^9,10^. Do the same hold true for canine cancers? A recent study reports similar mutation rates for mucosal melanoma between the two species^11^, while another study reports that canine and pediatric glioma are closer in mutation rate than canine and adult glioma^12^.

To answer these questions, we performed a pan-cancer and pan-breed study with 710 canine cases with public whole exome sequencing (WES) and/or whole genome sequencing (WGS) data^11–22^. We used 601 cases with WES data^11–13,15–17^ for discovery, and 18 with WES data^18,23^ and 91 with WGS data^18–22^ for validation. We also compared our canine findings to those published for the corresponding human cancers^3,4,24–27^. The study is described below.

## Results

### We performed a rigorous quality control (QC) on the WES data published for 601 canine cases

The 601 canine cases consist of over six cancer types, including mammary cancer^13^, oral melanoma^11^, osteosarcoma^15^, lymphoma^16^, glioma^12^ and hemangiosarcoma^17^ (Table S1). They represent over 35 breeds (Table S1A). Their WES data were generated by different groups, using different exome-capturing kits and Illumina sequencing machines^11–13,15–17^. We hence performed a rigorous QC to ensure that each study met a set of quality standards before any further integrative analysis.

Among these cancers, the mammary cancer study^13^ has the most comprehensive case information provided in the SRA database and related publications^11–13,15–17^ (Table S1). It has not only the patient info (e.g., age, sex, breed), but also histological subtypes and limited clinic data (e.g., tumor invasiveness, patient alive/death status). The osteosarcoma^15^, lymphoma^16^, glioma^12^, and hemangiosarcoma^17^ studies all have patient info (Table S1), but lack clinical data, while the melanoma study^11^ lacks this patient info, including the breed.

For our WES data QC, we first found that, except for the normal samples in the mammary cancer dataset, all other datasets have a median sequencing amount of >50 million (M) read pairs (Figure S1A). We identified two outliers with sequencing pairs <5M and excluded them from further analyses (Table S1B).

We then investigated the mapping of read pairs to the canine reference genome^28^. With the exception of the glioma study^12^, the mapping rates of all other datasets have >80% read pairs in each sample that are coordinately and uniquely aligned to the reference genome, with the median close to or above 90% (Figure S1B). We excluded 6 samples with mapping rates <60% from further analysis (Table S1B). Furthermore, all samples have >70% of their reads with mapping quality scores >30, with the mammary^13^ and melanoma^11^ datasets reaching median values of >90% (Figure S1B). For the target mapping rate, all studies have on average >50% uniquely and coordinately mapped pairs aligned to the coding sequence (CDS) regions, with the melanoma study^11^ achieving >60% (Figure 1A). We excluded three samples whose target mapping rate <30% from further analysis (Table S1B). All studies have reached an average CDS region read coverage median value of >70X among their samples (Figure 1B). We excluded 14 samples with coverage <30X (Table S1B). For the mapped read distribution in the target regions (which reflects sequencing randomness), we determined the deviation of each sample from its theoretical Poisson distribution (as a completely random sequencing process can be approximated by the Poisson distribution). The results indicate that the mammary cancer study has the most random sequencing, closely followed by the melanoma dataset (Figure 1C). Finally, for total callable bases, determined from MuTect^29^ (see later), of >10Mb (Figure 1D)all but one sample have >10Mb (Figure 1D). We excluded that sample.

**Figure 1.**
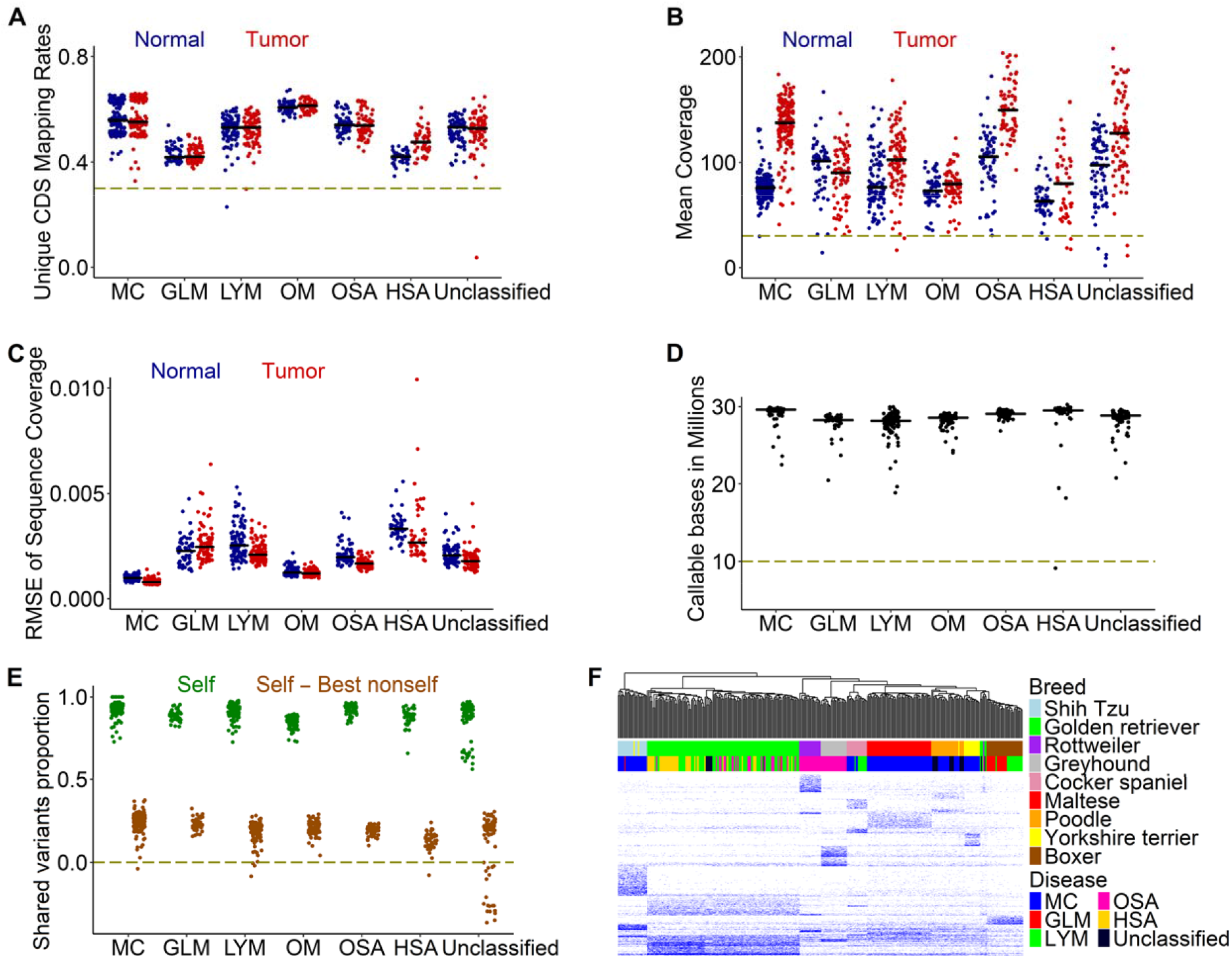
We performed a rigorous quality control (QC) of whole-exome sequencing (WES) data published for 607 canine cases. A. Mapping rates of read pairs aligned coordinately and uniquely to the canFam3 coding sequence (CDS) regions per sample. The median mapping rate of each dataset is indicated by a black line. The dashed line indicates the QC cutoff. MC: mammary cancer; GLM: glioma; LYM: lymphoma; OM: oral melanoma; OSA: osteosarcoma; HSA: hemangiosarcoma. B. Mean read coverage per sample. The figure is presented as in **A**. **C.** Root-mean-square error (RMSE) between the actual sequence coverage and the theoretical coverage based on the Poisson distribution per sample. The figure is presented as in **A.** D. Total number of callable bases per cancer case, output by Mutect (v 1.1.7). E. Tumor-normal pairing accuracy. “Self” (in green) is the proportion of germline variants shared between the normal and tumor samples of a dog. “Best nonself” is the proportion of germline variants shared between a normal or tumor sample of one dog and its best matched sample from another dog. “Self – Best nonself” (in red) indicates the difference, and a negative difference points to incorrect tumor-normal pairing. F. Breed validation with breed-specific germline variants. The heatmap indicates the variant allele frequency (VAF) values of each of 7,168 breed-specific variants in 359 total normal samples. See also Figure S1 and Table S1.

To assess the data accuracy, we examined the reported tumor and normal sample pairing using germline variants found in each sample (Figure 1E), assuming that correctly paired samples, compared to other samples in the same study, should share the most variants. We found a total of 24 mispaired cases and excluded them from further analysis.

Finally, we assessed the accuracy of the provided breed data. We first identified 7,168 breed-specific variants, which are germline variants that are unique to or enriched in each of the 9 breeds with ≥10 animals (see Materials and Methods). We then examined the variant allele frequency (VAF) of each breed-specific variant in normal samples (Figure 1F), and we identified 9 cases with mis-assigned pure breeds (5 Golden Retrievers, 3 Yorkshire Terriers and 1 Poodle), and excluded them from any breed-related analysis. We also repeated this analysis by including 107 cases with missing breed data (e.g., the oral melanoma dataset). We were able to unambiguously assign breeds to 46 cases (8 to Boxer, 10 to Cocker Spaniel, 14 to Golden Retriever, 4 to Maltese, 6 to Rottweiler, 3 to Shih Tzu and 1 to Yorkshire Terrier) (Figure S1D; Table S1F).

In summary, our QC analysis indicates that the mammary cancer and melanoma studies have the highest sequence quality, and that the mammary cancer study has the most comprehensive case info. A total of 50 cases out of 601 total have failed our QC steps (Table S1B) and are excluded from the analyses described below.

### Each cancer type has distinct molecular features

With 557 tumors from 551 cases (5 oral melanoma cases have both primary and metastatic tumors) that have passed our QC measures, we identified mutated and/or amplified/deleted genes, along with their molecular pathway, in each tumor. The study reveals unique alteration features for each canine cancer type.

Mammary cancer harbors frequent PI3K pathway alteration (54%) (Figure 2A). The PIK3CA H1047R mutation^30^ is especially common, found in 39% tumors (Figure 2A). However, another PIK3CA mutation hotspot, the E542/545 site, is intriguingly missing in these canine tumors, unlike human breast cancer^31^.

**Figure 2.**
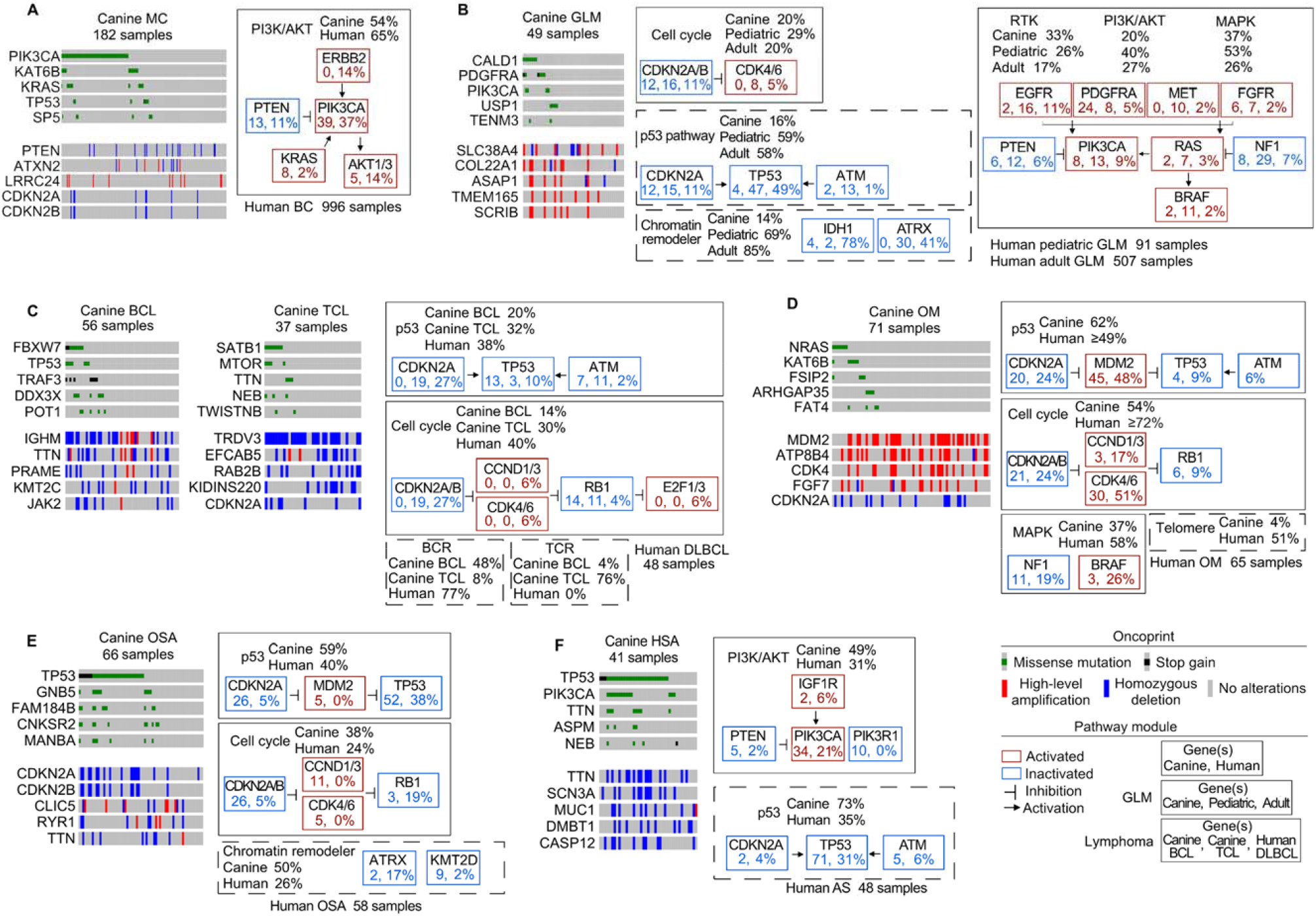
Each canine cancer type harbors distinct molecular features, many of which match those of their human counterpart. A-F. Oncoprints (left) indicate top 5 most frequently mutated genes and top 5 most frequently amplified or deleted genes in MC, GLM, B-cell lymphoma (BCL), T-cell lymphoma (TCL), OM, OSA and HSA. Shown on the right for each subfigure are the most frequently altered pathways reported in the corresponding human cancer, with genes mutated and/or amplified/deleted in at least 5% of either human or canine tumors indicated. The total percent of samples altered in each species per pathway is also shown. Pathways altered at a comparable frequency between the two species are surrounded by solid lines, while pathways where the alteration frequency between the two species differs by two-fold or greater are surrounded by dashed lines. See the bottom right corner for other legends. Human breast cancer (BC), pediatric and adult glioma (GLM), diffuse large B-cell lymphoma (DLBCL), oral melanoma (OM), osteosarcoma (OSA), and angiosarcoma (AS) are from published studies^3,4,24–27^. See also Figure S2 and Table S1B.

Oral melanoma and osteosarcoma both harbor frequent p53 pathway alteration (62%) (Figures 2D and 2E). However, the actual altered genes differ, with *TP53* mutated in 54% of osteosarcoma tumors (Figure 2E) and *MDM2* amplified in 48% of oral melanoma tumors (Figure 2D), consistent with previous findings^11,15^. Moreover, while deletion is more common in osteosarcoma, amplification is more common in oral melanoma (Figure S2). Indeed, *CDKN2A* is deleted in 26% of osteosarcoma and *CDK4* is amplified in 30% oral melanoma, resulting in frequent cell cycle gene alteration in both cancer types (Figures 2D and 2E).

Hemangiosarcoma has a TP53 mutation frequency of 71%, the highest among the 6 cancer types (Figure 2F). *PIK3CA* is another frequently mutated gene, found in 34% hemangiosarcomas. Consistent with published studies ^11,13,15,16^, T cell lymphoma harbors frequent T cell receptor (TCR), while B cell lymphoma harbors frequent B cell receptor (BCR) alteration (Figure 2C).

We noted that each canine cancer type shares major pathway alterations with its human counterpart (Figure 2), including breast cancer, pediatric and adult glioma, diffuse large B-cell lymphoma, oral melanoma, osteosarcoma, or angiosarcoma^3,4,24–27^. In both species, p53 pathway alteration and cell cycle alteration are common in osteosarcoma and oral melanoma (Figures 2D and 2E). PI3K signaling is the most frequently altered pathway in mammary cancer in both humans and dogs (Figure 2A).

We also observed dog-human differences. For example, p53 pathway alteration is at least twice more common in human glioma than in canine glioma (Figure 2B). Chromatin remodelers are frequently altered in human glioma but not in canine glioma (Figure 2B), while the opposite is observed for osteosarcoma (Figure 2E).

### Canine mutation rate varies among cancer types but not among breeds

We investigated mutation rate in each of the 557 canine cancers that pass the QC. Resembling human cancer^2,32^, canine mutation rate varies among these cancers, ranging from 0 to 161 somatic mutations per Mb CDS (Figure 3). However, overall the mutation rate is low, with a median of 0.6. Hypermutation (mutation rate >10) and ultrahypermutation (mutation rate >100) are both rare in these canine cancers (Figure 3).

**Figure 3.**
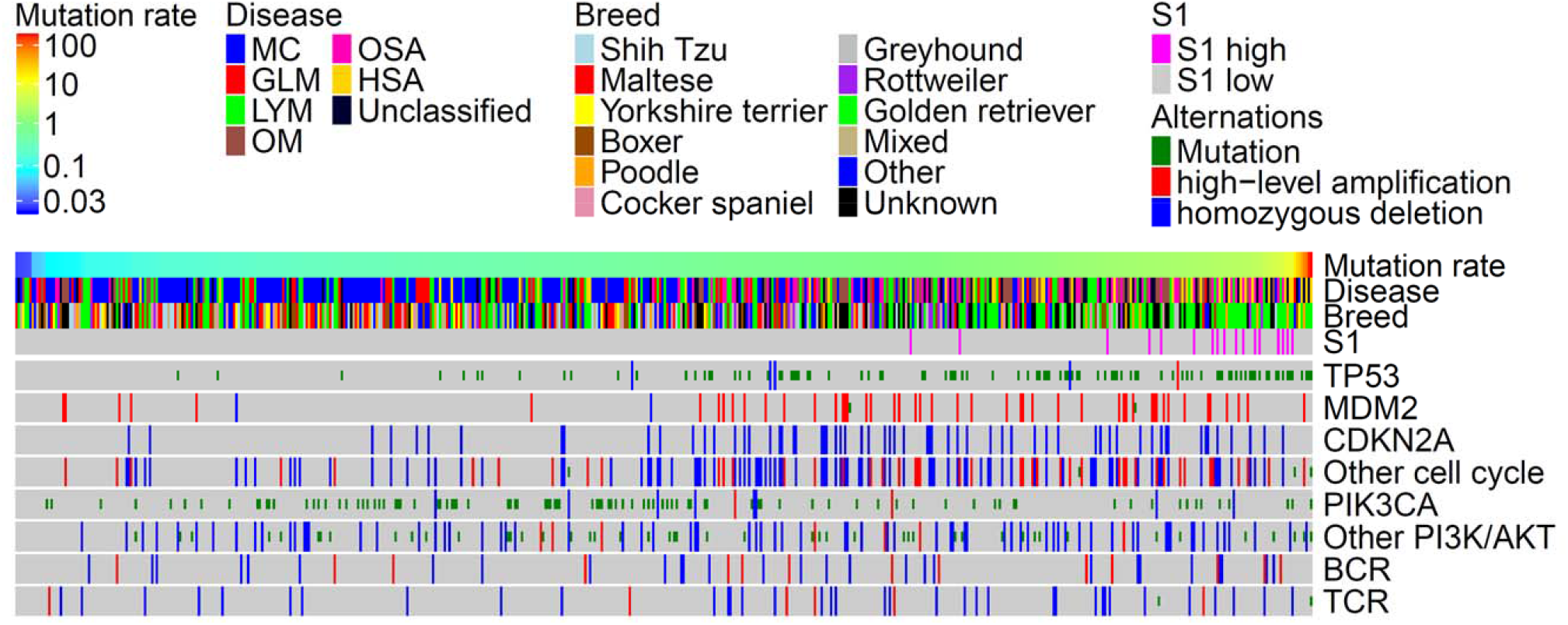
We identified mutation rate and alteration in each of 557 tumors representing over 6 cancer types and over 35 breeds. The oncoprint indicates mutation rate, cancer type, breed, and major gene/pathway alterations in 557 cancer cases (Figure 1). Cancers are ordered from left to right by lowest to highest mutation rate. Six cancer types (182 MC, 49 GLM, 91 LYM, 71 OM, 65 OSA, and 41 HSA tumors) as well as those with unknown cancer types (55 cases) are included. Specific breeds shown include those validated (350 cases; Figure 1) or clearly assigned by our analysis (46 cases; Figure S1). Also shown include mixed breeds (21 cases), other breeds (61 cases) for which we could not validate due to small sample size, and unknown (76 tumors for 70 cases) where breeds were neither reported nor assigned by us, or breeds that failed our validation (9 cases). See also Figure S3 and Table S3.

The mutation rate varies among cancer types (Figure 4A). Canine mammary cancer and glioma have the lowest mutation rate, with a median value of 0.37 and 0.43 respectively, and hence are classified as mutation rate-low (MR-L) (Figure 4A). Canine oral melanoma, osteosarcoma, and hemangiosarcoma all have higher mutation rate, however, with a median value of 0.94, 1.07 and 1.28 respectively. They are hence classified as MR-high (MR-H) (Figure 4A). Canine lymphoma is somewhere in between, with a median of 0.6, and is classified as MR-median (MR-M) (Figure 4A). These findings are validated using two other WES datasets^18,23^.

**Figure 4.**
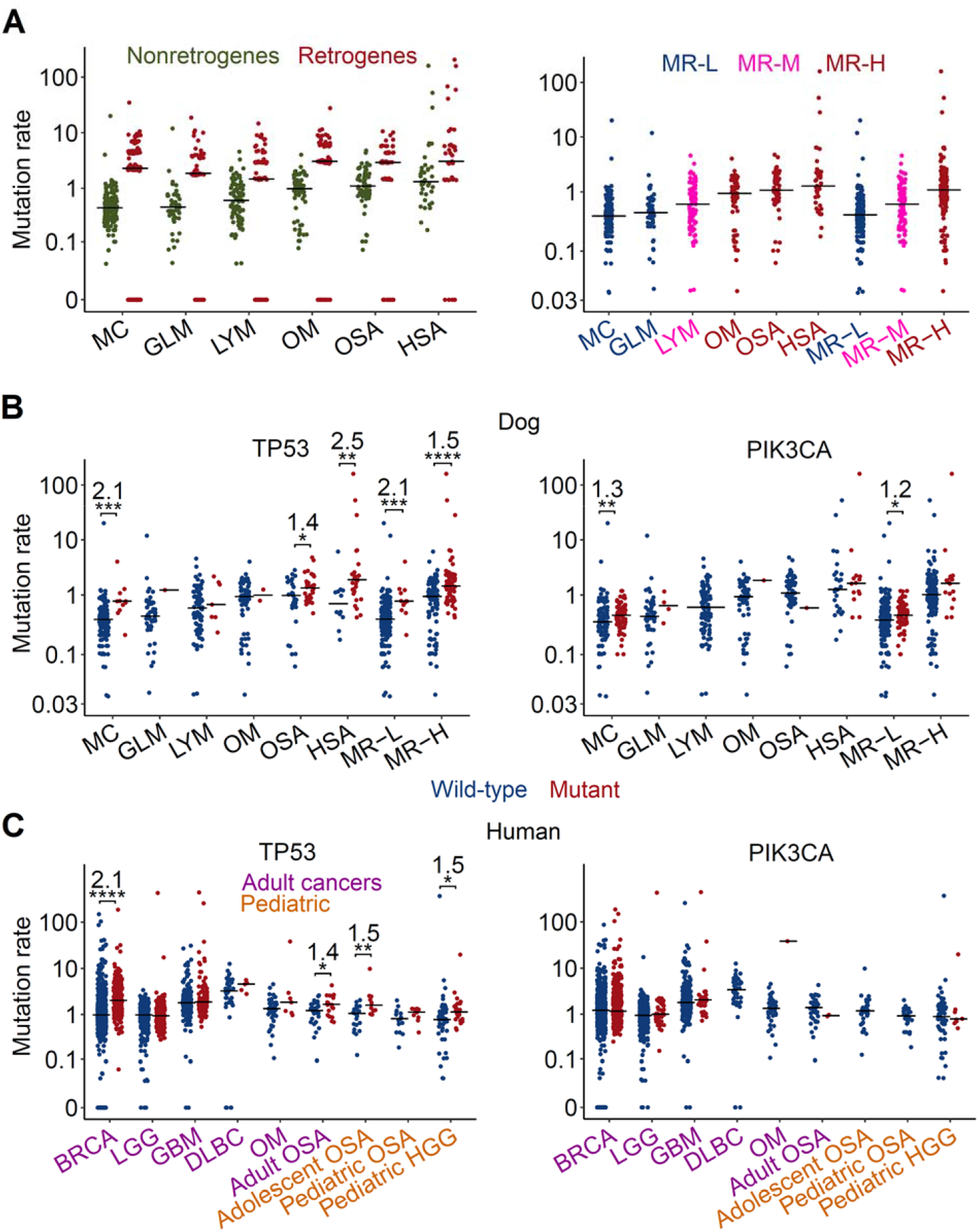
Mutation rate varies across cancer types and is correlated with TP53 mutation. A. Mutation rate distributions for each cancer type, ordered (left to right) from lowest to highest median values. The left plot shows that retrogenes, a type of pseudogene, have significantly higher mutation rates than non-retrogenes, and are excluded from further analyses. The right plot indicates that canine tumors are classified into mutation rate-low (MR-L), -medium (MR-M) and -high (MR-H). B&C. Mutation rate distributions for samples of each cancer type with (Mutant) or without (Wild-type) TP53 (left) or PIK3CA (right) mutations in dog (B) and human (C) cancers. Wilcoxon tests were conducted if both Wild-type and Mutant groups per cancer contained >= 3 cases to identify those cancer types with a significant association between TP53 or PI3KCA mutation and tumor mutation rate. *, **, *** and **** represent p<0.05, <0.01, <0.001, and <0.0001 respectively. For cancer types with a significant association found, the fold change in median mutation rate of TP53 (or PIK3CA) Mutant versus Wild-type samples is also shown. See also Figure S4 and Table S4.

Within the same cancer type, the mutation rate appears similar among breeds (Figure 5A), except for osteosarcoma, where Golden Retrievers have significantly higher mutation rates than those of Rottweilers and Greyhounds (Figure 5A). Thus, we conclude that mutation rates in canine cancers are primarily determined by cancer types, but not by breeds.

**Figure 5.**
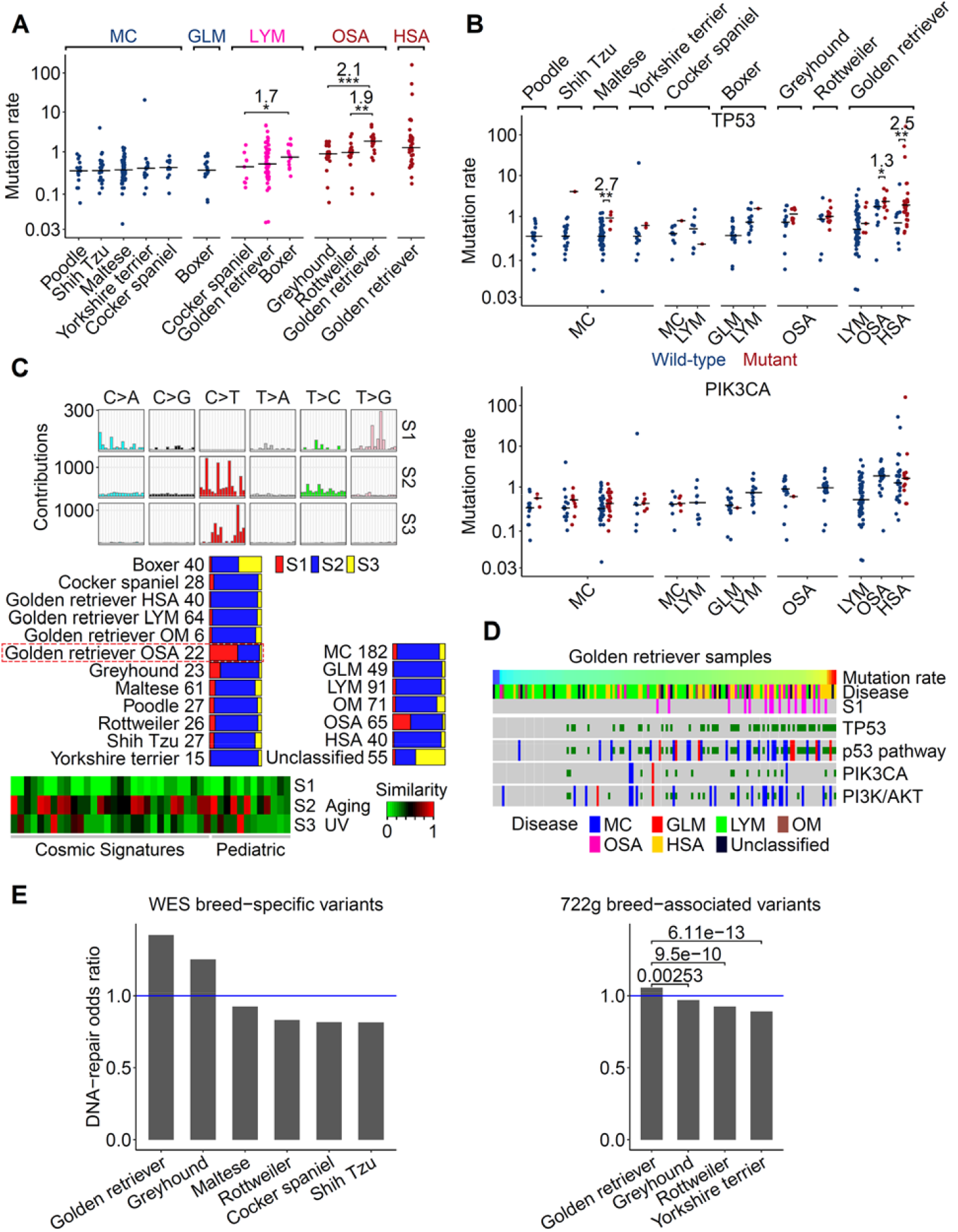
Mutation rate is largely independent of breed, and Golden Retrievers harbor distinct molecular features. A. Mutation rate distributions of cases grouped by cancer type and breed. Only groups with ≥ 10 cases are shown. B. Mutation rate distributions of cases grouped by breed, cancer type, and finally TP53 (top) or PIK3CA (bottom) mutation status. Only those with TP53 (or PIK3CA) Wild-type and Mutant combined cases of >10 are shown. The same statistical analysis is shown as described in Figure 4B. C. Three mutation signatures are detected in 553 canine tumors, after excluding one HSA outlier (top). The bar plot (middle) indicates the distribution of the three signatures in each cancer type and in each validated breed. The numbers denote the tumor sample counts. The bottom heatmap indicates the cosine similarity scores between each canine signature and each of the 30 COSMIC ^32^ and 12 pediatric ^3,4^ signatures. D. Golden Retriever-specific oncoprint, including 149 animals, presented as described in Figure 3. E. Golden Retrievers have the most variance in DNA repair genes among all breeds examined. The left bar plot indicates the odds ratios of breed-specific variants (see Figure 1) inside DNA-repair genes (116 total) versus non-DNA-repair genes, normalized by total CDS base pairs for the corresponding gene types. The right plot indicates the same analysis with all germline variants (breed-specific and -nonspecific) in each breed reported by a published study^59^. The blue line denotes the odds ratio where the variants distribution is balanced between the two gene types. *P*-values are from Fisher’s exact tests. See also Figure S5 and Table S5.

### Canine mutation rate is correlated with TP53 mutation but not PIK3CA mutation

Like human cancer, TP53 is frequently mutated in canine cancer (Figure 3). Importantly, we noted a strong association between mutation rate and TP53 mutation across cancer type and within a cancer type (Figure 4B). This is clearly seen in canine hemangiosarcoma and osteosarcoma, which are both MR-H (Figure 4A) and have TP53 mutated in 68% and 58% of tumors, respectively (Figure 4B). Furthermore, the median mutation rate in osteosarcomas and hemangiosarcomas with mutant TP53 ihas increased to 1.34 and 1.88 respectively, compared to 0.98 and 0.7 for the corresponding tumors with wild type TP53 (Figure 4B).

A strong association between mutation rate and TP53 mutation is also observed across breeds (Figure 5B). Indeed, the median mutation rate increases significantly with TP53 mutation in Golden Retriever (0.66 to 1.88), Greyhound (0.73 to 1.14), Rottweiler (0.68 to 1.0) and Maltese (0.34 to 0.91) (Figure 5B).

PIK3CA is another gene frequently mutated in canine cancer (Figure 3). However, in contrast to TP53, we did not observe an association between mutation rate and PIK3CA mutation in any cancer type or breed (Figures 4B and 5B). Furthermore, for BCR and TCR alterations, which are common only in lymphoma (Figure 2C), we observed an association only for TCR alterations (Figure S4) but not for BCR alterations.

Mutation rate is consistently associated with p53 pathway alteration (Figure 3). This is because besides TP53, we also observed a strong association with other genes in the p53 pathway, including CDKN2A deletion which is frequent in canine osteosarcoma and melanoma (Figures 3 and S4), and ATM mutation which is frequent in lymphoma (Figure S4).

Cell cycle pathway alteration is also associated with canine mutation rate (Figure 3), due to CDKN2A/2B deletion and CDK4 amplification (Figure S4).

Notably, we observe the same associations described above about canine cancers in corresponding human cancers, either adult or pediatric (Figure 4B). Our findings add another piece of data supporting that canine cancers faithfully recapitulate the molecular features of their human counterparts.

### Osteosarcoma in Golden Retriever harbors mutation rate-associated signature

We investigated mutation signatures in these canine tumors, and identified three signatures, which we named S1, S2 and S3 (Figure 5C). S2, matching the aging signature reported in human adult and pediatric cancers^2–4,32^, is the dominant signature in these canine cancers (Figure 5C). S3 matches the human UV signature^2–4,32^, and is mostly enriched in tumors of unknown cancer types (Figure 5C). S1 lacks significant matches to any known signatures reported in human cancer (Figure 5C) ^2–4,32^. Notably, S1 is significantly enriched only in osteosarcomas of Golden Retriever dogs (Figure 5C); hence it is considered breed- and cancer-specific. S1 is associated with mutation rate, but not with TP53 mutation (Figures S5A-B). We also detected S1 by analyzing published WGS data of canine cancer^18–22^.

To better understand the genetic basis of S1, we determined S1-associated mutations, which include ABTB2 R643C, a germline mutation limited to Golden Retrievers within our study, and BRPF1 V771G, a somatic mutation only found in osteosarcoma of Golden Retrievers (6 total) (Figure S5C). Notably, BRPF1 encodes a subunit that constitutes the MOZ/MORF histone acetyltransferase complexes, which remodel chromatin, regulate gene expression and is implicated in cancer development^33^.

### DNA repair genes appear to evolve faster in Golden Retrievers

Compared to other breeds, Golden Retriever dogs harbor higher mutation rates, more TP53 mutation, and a unique mutation signature (Figures 5A-C). To understand the reasons for these observations, we examined DNA repair genes, whose alterations lead to inefficient DNA repair in the cell and higher mutation rates^2,34^. We determined the ratio of germline mutations in classical DNA repair genes (116 total; see Table S5G) versus those in all non-DNA repair genes in the canine genome. We found that Golden Retriever has the highest ratio, compared to other breeds examined, in two datasets including our WES data described above and a published WGS study of 722 dogs^35^ (Figure 5E). The results indicate that DNA repair genes evolve faster in Golden Retriever, which may decrease DNA repair efficacy and possibly explain the observed higher mutation rates in Golden Retriever dogs (Figure 5A).

## Discussion

Taking advantage of public canine data, we have investigated 601 canine cases of over 6 cancer types and over 35 breeds. To our knowledge, this represents the first pan-cancer and pan-breed study for the dog. Importantly, our study provides a deeper understanding of canine cancer mutation rates.

### Mutation rate and TP53 mutation

Our study reveals that canine mutation rate is primarily cancer type-dependent, but largely breed-independent. TP53 mutation and p53 pathway alteration are potential reasons, due to their strong association with mutation rate observed across cancer types and across breeds, a pattern not observed in PIK3CA mutation.

Canine melanoma, osteosarcoma, and hemangiosarcoma have significantly higher mutation rates than other cancer types investigated. Notably, TP53 mutations are very frequent in canine osteosarcoma and hemangiosarcoma, while MDM2 amplification (which promotes TP53 degradation) is common in melanoma. In contrast to these cancers, canine mammary cancer and glioma harbor infrequent TP53 mutation or p53 pathway alteration, and a lower mutation rate. We hypothesize that these observations are related to the cells of origin in each cancer and their tumorigenesis mechanism.

Mammary cancer originates from epithelial cells. Establishment of epithelial cell apical-basolateral polarity decreases cell proliferation, and acts as potent tumor suppressor^36,37^. PIK3CA H1407R/K mutation, common in canine mammary cancer, increases cell stemness^38^ and decreases epithelial cell polarity, leading to accelerated cell proliferation and tumorigenesis. For glioma, epigenetic alterations (e.g., aberrant DNA methylation^12^) drives tumorigenesis. However, in both cancer types, even with accelerated cell proliferation, the cell cycle checkpoint is functional, leading to low mutation rate.

Hemangiosarcoma, osteosarcoma and melanoma all arise from mesenchymal cells, which lack cell polarity and cell adhesion. Loss of function of p53, due to either TP53 mutation or MDM2 amplification, leads to defective cell cycle checkpoints. This shortens the G0 and G1 phases, accelerating cell cycle. As a result, fewer DNA damages are repaired and fewer DNA replication errors are corrected^9^, increasing the mutation rate and contributing to tumorigenesis.

In supporting the hypothesis above, we have noted a strong association between cell cycle alteration, including CDK2A deletion, and canine mutation rate.

### Dog-human homology

Our pan-cancer study reveals numerous dog-human homologies. First, each canine cancer type shares major pathway alterations and often the same mutated genes with its human counterpart, consistent with previous individual cancer studies^11–22^. Second, the order of canine cancer types sorted by the mutation rate (i.e., mammary cancer < glioma < lymphoma, etc.) is the same as that of the corresponding human cancer types^11,39–43^, even though the actual mutation rates may differ. Third, across cancer types in both species, mutation rate is strongly associated with TP53 mutation and p53 pathway alteration, but not with PIK3CA mutation.

### Golden Retriever

Among the 35 breeds investigated here, Golden Retriever is the largest breed, with 149 animals in total after QC and breed validation. These dogs constitute a large portion of the osteosarcoma and lymphoma cases and the entirety of the hemangiosarcoma case set. Importantly, our study reveals unique features of Golden Retrievers. These include a mutation signature, which is enriched only in osteosarcomas of Golden Retriever dogs and is associated with mutation rate, independent of TP53 mutation. Furthermore, DNA repair genes, which are closely linked to mutation rate^8,9^, appear to evolve faster in Golden Retrievers than other breeds examined. This may lead to less efficient DNA repair, possibly explaining the higher mutation rate of cancers from Golden Retrievers. Importantly, this may explain why Golden Retriever dogs are prone to the development of multiple cancer types. These of course require future validation studies.

## Materials and Methods

### WES data collection

We acquired canine cancer data from the Sequence Read Archive (SRA) database with BioProject IDs PRJNA391455 (osteosarcoma), PRJNA489159 (mammary cancer), PRJNA247493 (lymphoma), PRJEB12081 (oral melanoma), PRJNA579792 (glioma), PRJNA552034 (hemangiosarcoma), and PRJNA247493 (unclassified). We also obtained other information from relevant publications^11–17,30,44,45^. Sources are shown in Table S1.

### WES data QC

#### Mapping rates

We aligned sequence read pairs using BWA-aln (version 07.17)^46^ to the canine reference genome CanFam3.1. We identified coordinately and uniquely mapped pairs based on the flag values and TAG values (with XT: AU or XT: AM) to calculate each sample’s overall mapping rate. We identified pairs with at least one read with >=1bp overlap with a coding sequence (CDS) region (from Canfam3 1.99 GTF) to calculate the target mapped rate. A case was considered to have failed QC if its tumor or normal sample has a coordinately and uniquely mapping rate below 0.6 or a target mapping rate below 0.3.

#### Mapped sequence coverage and distribution

We calculated base coverage using GATK (version 3.8.1)^47^ DepthOfCoverage with minimum mapping quality and base quality scores set to Phred 10. The mean coverage was derived by dividing the sum of the total depth for each locus by the number of total loci. Cases were considered to have failed QC if either their tumor or normal samples have a mean coverage below 30X.

To identify the read distribution in target regions (which reflects sequencing randomness), we calculated the root mean square error (RMSE) of sequence coverage by comparing the sequence distribution in the sample to Poisson distribution with λ equals to the mean coverage of each sample.

#### Quantifying Callable bases

We calculated callable bases for each case using Mutect (v1.1.7)^29^ with minimum base quality score set to Phred 30. Cases with fewer than 10 million callable bases were ruled out before any integrative data analysis.

### Germline variant calling

Starting with realigned bam files (see Somatic mutation detection in WES data), we used GATK 3.8-1^47^ HaplotypeCaller to call germline variants for every normal and tumor sample separately (parameters: -dontUseSoftClippedBases -stand_call_conf 20.0). We then filtered out variants using GATK VariantFiltration (filters: FS > 30.0 and QD < 2.0). We calculated the coverage of every coding sequence locus using GATK DepthForCoverage with parameters --minBaseQuality 10 and --minMappingQuality 10. Finally, we compiled all protein modifying variants (indels, stop-gain, and missense) that are observed in at least one case and called in both of its normal and tumor samples. We calculated the Variant Allele Frequency (VAF) for each called variant in every sample using the formula: allele depth/total base depth. For variants that are not called by HaplotypeCaller, we assigned a VAF value of 0. For each variant, we excluded samples with locus coverage <10, according to DepthForCoverage tool, from further analysis related to that variant, whether it has been called or not.

### Tumor-normal sample pairing accuracy

Using all passed germline variants by VariantFiltration above, we calculated the total number of variants in tumor (*T_i_*) and normal (*N_i_*) sample for every case *i*. We also calculated the number of shared variants between any two tumor and normal samples *S_i,j_* from the same dataset. The proportion of shared variants between a tumor sample *i* and a normal sample *j* is given by:

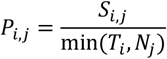

I.e. it is the number of shared variants divided by the smaller number of variants amongst two samples. If *i* = *j*, *P_i,j_* = *P_j,i_* = *P_i,i_* = *Self_i_*, the proportion of shared variants between the tumor-normal pair of the same case. The *Best nonself_i_* match is the highest proportion of shared variants observed between the tumor sample of case *i* and any other normal in the same dataset or between the normal sample of case *i* and any other tumor, whichever is higher.

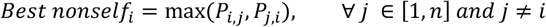

Therefore, *Self_i_* – *Best nonself_i_* is negative if and only if either the tumor or the normal of a case *i* has a better match from another case.

### Breed validation

Using the compiled germline variants above, we analyzed the VAF values of all normal samples. Each breed with at least 10 samples was included and considered a major breed group in the analysis. All samples with assigned breed labels other than major breeds were grouped into “Other” breed. All samples of mixed or missing breed were excluded from this analysis.

We categorized VAF values into reference (VAF < 0.2) and non-reference (VAF ≥ 0.2), or NA for variants with locus coverage below 10. In this analysis, we first filtered out variants whose locus coverage <10 in more than 20% of the total samples. This is needed to rule out variants with poor coverage or not covered in every whole-exome library.

We considered a variant “breed-specific” if it falls into breed-unique or breed-enriched variant category. A breed-unique (for a major breed A) variant satisfies three criteria: 1) is nonreference in at least 5 samples of breed A; 2) is non-reference in at least 40% of breed A samples; and 3) is reference in all other samples with sufficient coverage. Breed-enriched variants must be enriched in one and exactly one breed. For each variant, we used Fisher’s exact test to compare each combination of two breeds using the reference and non-reference sample counts in each breed. A breed-enriched variant must be 1) enriched in breed A against every other breed group at *P*=0.1 and 2) not enriched in any breed B against any breed group at *P*=0.1.

Starting with all breed-specific variants found above, we omitted any sample with low coverage in more than 20% of the breed-specific variants because these samples can introduce noise in the clustering process. We assigned random VAF values for the remaining low coverage sites. We finally performed standard hierarchical clustering. Cases of major breeds with breed label and breed cluster agreement pass the QC. We also assigned breeds to cases with missing breed info based on their breed cluster (Table S1E-F).

### Identification of somatic copy number alterations

We used VarScan v2.4.2^48^ to call somatic copy number alterations (CNAs) using WES data of matched tumor and normal sample pairs. We applied the software CBS^49^ to coding regions of the identified CNAs via DNAcopy R package^48^. Segments with log2 (T/N) greater than 1 or less than −1 were regarded as candidate CNAs. For canine MC samples, candidate CNAs from CBS results were further selected if log2 (T/N) associated with a gene is positively correlated with its mRNA expression (correlation coefficient > 0.2 for both Spearman and Pearson correlations). For other canine cancers analyzed, we selected genes with CNAs detected by both CBS and SEG^50^ using the same cutoff as mentioned^50^.

### Comparison of genomic alterations between canine and human cancer at pathway level

Mutated genes and altered pathways in human cancers were extracted from published studies, including 996 breast cancers^24,25^, 91 high grade pediatric gliomas^40,41^, 507 adult low grade gliomas^12^, 48 diffuse large B-cell lymphomas^25^, 65 oral melanomas^27^, 58 pediatric osteosarcomas^3,4^, and 48 angiosarcomas^26^.

### Somatic mutation detection in WES data

To acquire the tumor mutation burden, we mapped samples with BWA (version 0.7.17)^46^ and used Picard (version 2.16.0) to exclude unmapped reads, sort, and mark duplicate sequences. We then used GATK (version 3.8-1)^47^ to realign the samples and find exonic bases with sufficient coverage. We then used Mutect (version 1.1.7)^29^ to detect somatic mutations with reference genome CanFam3, with a minimum base quality of 30 ^51^. We applied further filtering as previously described ^11^. We detected somatic indels with Strelka, and further filtered for those found in the CDS region of CanFam3^52^. Finally, we annotated mutations for function with Annovar (version 2017Jul16)^53^, and separated retrogenes from known coding genes for mutation rate calculation. Additionally, using the realigned bam files from GATK, we used the GATK commands HaplotypeCaller and VariantFiltration to detect the germline mutations present in the WES data. These germline mutations were then also annotated with Annovar. The number of mutations in coding nonretrogenes of passing sequence quality by Mutect was divided by the number of callable exonic bases to get the mutation rates.

### Identifying retro and nonretrogenes

We acquired the retrogene and nonretrogene lists from the Canfam3 1.99 GTF file. We classified a gene as a retrogene if it has an Ensembl ID but no gene name, the gene is protein-coding and it contains only one exon. A nonretrogene has both an Ensembl ID and a gene name, the gene is in the protein-coding region, and it contains more than one exon. We also excluded mitochondrial genes to get the final retrogene and nonretrogene list, leaving 1564 retrogenes and 14733 nonretrogenes in the list.

### Quantifying the Consensus Somatic Mutation Calling

We used GATK4 Mutect2 (version4.1.6) ^41^,Varscan2 (version2.4.2)^54^, and LoFreq^55^ (version2.1.2) to call somatic variants. We first ran Mutect2 in panel-of-normals (PON) mode using paired normal samples that pass our QC (n=557). Using the resulting PON file, we used Mutect2 to call somatic variants in paired mode with parameters: –germline-resource: DbSNP_canFam3_version151-DogSD_Feb2020_V4.vcf (SNP derived from Ensembl (version 151), DogSD germline SNP (downloaded from ftp://download.big.ac.cn/idog/dogsd/vcf/Filtred_Published.vcf.bz2, and Genome-Wide Variant Discovery^56^) –af-of-alleles-not-in-resource 0.008621. To find consensus variant callings across Mutect2, Varscan2, and LoFreq, We followed the procedure and parameters described ^22^ to identify the consensus somatic variants using SomaticSeq (version 3.4.1)^57^.

### Comparisons of TP53 and PIK3CA mutations on mutation rate

Mutation rate and alterations in the TP53, PIK3CA, and Cell Cycle pathways in cancers matching canine types (Breast invasive carcinoma (BRCA), Low grade glioma (LGG), Diffuse large B-cell lymphoma (DLBC), and Glioblastoma (GBM), all from the TCGA pan-cancer atlas) were acquired from cbioportal^27,41,42^. Oral melanoma (OM), osteosarcoma (OSA), and pediatric high-grade glioma (HGG) alterations/mutations for these pathways and genes were acquired from additional sources ^11,39,43^. Mutation rates between samples with any mutation in TP53 or PIK3CA or wild-type per cancer type are shown in Figure 3C. The difference between samples with mutated vs. wild-type TP53 and PIK3CA were compared by Wilcoxon rank-sum test, and the fold change between the medians of each group (altered/not altered) within cancers was shown if the difference was determined to be significant. Additionally, the same comparison was performed for samples with any mutation, gene fusion, or copy number change, which would be considered altered in this case, in any gene in these respective pathways, between altered and non-altered samples, as well as the full pathway (Supplemental Figure 4).

### Mutation signatures

The raw counts of filtered somatic mutations (see Somatic mutation detection in WES data) were used as input for signature discovery. We ran SignatureAnalyzer R code ^58^ with the default parameters, except for the number of iterations=40 and hyper=False. 553 total tumor samples were used after excluding one HSA ultrahypermutated sample, HSA_4. The most frequent solution (20 iterations) consists of four signatures, where the classic aging signature is broken into C>T and T>C signatures, with the latter showing no resemblance to any known signature. We considered this separation an artifact. The second most frequent solution (15 iterations) consists of three signatures where the classic aging signature has both C>T and T>C mutations. We selected this solution for the remaining analysis. The signature plots are based on raw mutation counts. To compare canine signatures to Cosmic signatures v2^32^, and pediatric cancer signatures^3,4^, we adjusted the human signatures using the human genomic background trinucleotide probabilities and the canine signatures using the canine exonic trinucleotide probabilities. We used cosine similarity to compare the adjusted signatures. All data related to this analysis are in Table S5.

### S1 association with germline variants and somatic mutations

We categorized Golden Retriever tumors into S1 high (N=16), containing 20 or more S1 mutations, and S1 low tumors (N=119), containing fewer than 20 S1 mutations. For germline variants, we started with the combined list of protein modifying variants (see Germline variant calling), which consists of 75,253 variants. Based on the VAF distribution analysis, we categorized VAF values into reference (VAF < 0.2), heterozygous (0.2 ≤ VAF < 0.8) and homozygous (VAF ≥ 0.8). We conducted two independent tests for every germline variant: (1) homozygous versus reference and (2) heterozygous versus reference.

Since S1 is Golden Retriever enriched, we controlled for breed by including only Golden Retriever specific variants. These are variants enriched in every Fisher’s test between Golden Retriever and another major breed at *P*=0.05. We found 514 homozygous and 730 heterozygous Golden Retriever-specific variants, some of which are overlapping. From the remaining list, we selected variants which are enriched in normal samples matching S1 high tumors versus those matching S1 low tumors at *P*=0.05. Only 17 heterozygous and one homozygous variants pass this condition. Finally, we used UCSC multiz100way protein alignment to select only evolutionarily conserved variants (Table S5F).

For somatic mutations, we started with a compiled list of 13,917 somatic mutations in 557 tumors (see Somatic mutation calling), but only 39 are observed in at least 5 samples, the minimum we required to perform enrichment analysis. We directly performed S1 high versus S1 low Fisher’s test on these 39 mutations and selected those enriched at False Discovery Rate 0.1. 13 mutations pass this condition. Among them, we selected the evolutionary conserved ones (Table S5F). Golden RetrieverGolden RetrieverGolden RetrieverGolden RetrieverGolden Retriever

### DNA-repair variants within breeds

We performed two independent analyses for DNA-repair variants: (1) We used the list of 7,168 breed-specific protein-modifying variants (see Breed validation) and counted the number of variants located inside DNA-repair and non-DNA-repair domains. The DNA-repair domain is defined as a list of 116 classical DNA-repair genes (Table S5G). (2) We used a list of 34 million genomic variants (SNPs or indels) inside gene regions identified using 722 WGS data of 722 dogs ^35^. In this dataset, our analysis was limited to breeds that have 10 or more samples and belong to one of the major breeds in this study. For each breed, breed-associated variants were calculated. Among the 34 million variants in the second set, a variant was considered breed-associated if it is called in at least half of that breed’s samples at a minimum base coverage of 10 reads and a minimum VAF value of 0.2.

In both data sets, the bar plots (Figure 5E) indicate the odds ratios of DNA-repair variants, which is the ratio between DNA-repair and non-DNA-repair variant counts divided by the ratio between DNA-repair and non-DNA-repair domain size. Finally, Fisher’s test was applied to test differences in the ratio of DNA-repair variants between two breeds using the DNA-repair and non-DNA-repair variant counts in the two breeds.

## Acknowledgments

We thank the Georgia Advance Computing Resource Center (GACRC) for supporting this work. This work is funded by NCI R01 CA182093.

